# The WY domain of an RxLR effector drives interactions with a host target phosphatase to mimic host regulatory proteins and promote *Phytophthora infestans* infection

**DOI:** 10.1101/2023.07.24.550319

**Authors:** Adam R. Bentham, Wei Wang, Franziska Trusch, Freya A. Varden, Paul R. J. Birch, Mark J. Banfield

**Affiliations:** Department of Biochemistry and Metabolism, John Innes Centre, Norwich Research Park, Norwich, NR4 7UH, UK; Department of Cell and Molecular Sciences, James Hutton Institute, Errol Road, Invergowrie, DD2 5DA, Dundee, UK; Division of Plant Sciences, College of Life Science, University of Dundee (at JHI), Errol Road, Invergowrie, DD2 5DA, Dundee, UK

**Keywords:** Potato, *Phytophthora infestans*, oomycete, plant-pathogen interactions, WY domain, protein-protein interactions, structural biology

## Abstract

Plant pathogens manipulate the cellular environment of the host to facilitate infection and colonization, often leading to plant diseases. To accomplish this, many specialized pathogens secrete virulence proteins called effectors into the host cell, which subvert processes such as immune signalling, gene transcription, and host metabolism. *Phytophthora infestans*, the causative agent of potato late blight, employs an expanded repertoire of RxLR effectors with WY domains to manipulate the host through direct interaction with protein targets. However, our understanding of the molecular mechanisms underlying the interactions between WY effectors and their host targets remains limited. In this study, we performed a structural and biophysical characterization of the *P. infestans* WY effector, Pi04314, in complex with the potato Protein Phosphatase 1-c (PP1c). We elucidate how Pi04314 uses a WY domain and a specialised C-terminal loop carrying a KVxF motif that interact with conserved surfaces on PP1c, known to be used by host regulatory proteins for guiding function. Through biophysical and in planta analyses, we demonstrate that Pi04314 WY or KVxF mutants lose their ability to bind PP1c. The loss of PP1c binding correlates with a reduced capacity to re-localize PP1c from the nucleolus and a decrease in lesion size in plant infection assays. This study provides insights into the manipulation of plant hosts by pathogens, revealing how effectors exploit key regulatory interfaces in host proteins to modify their function and facilitate disease.

## Introduction

During infection, plant pathogens manipulate host cells through the secretion of specialised proteins (effectors) with various functions to aid in colonisation of the host (Wirthmueller et al. 2013). These effectors act on many diverse pathways, functioning to subvert immunity, control host metabolic processes or modulate transcriptional regulation (He et al. 2020). While many host targets of plant pathogen effectors have been identified, there is very little structural or mechanistic data available describing how these interactions between pathogen and host proteins are mediated (Mukhi et al. 2020).

*Phytophthora infestans* is an oomycete pathogen responsible for late blight disease on a wide variety of solanaceous plants, resulting in significant losses to potato and tomato crops worldwide. *P. infestans* is predicted to deliver hundreds of effectors that manipulate host processes to facilitate infection and suppress plant immunity. Among these are the RxLR effectors, named for the conserved Arg-x-Leu-Arg (RxLR) motif at their N-terminus required for translocation into plant cells (Whisson et al. 2007). RxLR effectors target various host proteins and modulate their functions through direct interaction and a range of modes-of-action (Boevink et al. 2016; He et al. 2020; McLellan et al. 2022).

A subset of RxLR effectors contain one or several tandem WY domains at their C-terminus, which are important for host target specificity and effector function (Zhang et al. 2019; Li et al. 2023). WY domains are found in diverse oomycete species and are characterised by an 𝛼-helical fold with a buried hydrophobic core (Mukhi et al. 2020). Outside the hydrophobic core, WY effectors demonstrate high sequence variation, making it difficult to elucidate the function and evolution of the WY domain, but it has often been implicated in mediating protein-protein interactions (King et al. 2014; Zhang et al. 2019; Boevink et al. 2016). An additional class of WY effectors, called LWY effectors, carry a conserved leucine (L) residue prior to the canonical WY residues. This additional leucine is thought to be important for the structural organisation of tandem WY domains in LWY effectors, such as PSR2, creating joint-like structures that could allow the flexibility for conformational change upon interaction with other proteins, including host targets (Zhang et al. 2019; Li et al. 2023; He et al. 2019). Intriguingly, WY domains can be decorated with additional protein-interaction motifs to aid target selectivity. For example, the *P. infestans* RxLR effector PexRD54 comprises five WY domain tandem repeats and a C-terminal ATG8-interaction motif (AIM) that directly binds the potato Autophagy-related protein 8 (ATG8), outcompeting host autophagy receptors (Maqbool et al. 2016; Dagdas et al. 2016).

The RXLR effector Pi04314 has been shown to enhance *P. infestans* leaf colonization through interaction with the host Protein Phosphatase 1-c (PP1c) enzyme, potentially manipulating expression of jasmonic and salicylic acid-responsive genes in the host (Boevink et al. 2016). The Pi04314 effector is predicted to have a single WY domain, followed by a KVxF motif. The KVxF motif is a short peptide motif first discovered in the regulatory subunit MYPT1 (myosin phosphatase targeting subunit 1) that is important for interacting with the hydrophobic grooves on protein phosphatases, such as the PP1 proteins (Westphal et al. 1999; Nygren and Scott 2015; Terrak et al. 2004). Intriguingly, Pi04314 associates with three potato PP1 catalytic subunits (PP1c-1, PP1c-2 and PP1c-3) with the Pi04314 KVxF motif important for the function of the effector. This interaction causes the re-localization of PP1c isoforms from the nucleolus to the nucleoplasm, forming a holoenzyme that promotes susceptibility (Boevink et al. 2016).

In this study, we report the structural basis of Pi04314 interaction with PP1c, highlighting how the WY domain and KVxF motif bind key interfaces for PP1c function to facilitate *P. infestans* infection. Through structural, biophysical and in planta assays we describe the roles of the Pi04314 WY domain and KVxF motif in PP1c binding and show how these data correspond with *P. infestans* infection assays and Pi04314-mediated nuclear re-localisation of PP1c. Our findings provide insights into how an oomycete effector targets and manipulates a key host enzyme through conserved regulatory interfaces to promote disease.

## Results

### The crystal structure of Pi04314 bound to PP1c establishes the WY domain and KVxF motif as important for target binding

To better understand the molecular mechanisms underpinning Pi04314 function, we determined the crystal structure of the effector in complex with the host target PP1c isoform-3 (herein simply referred to as PP1c). A complex of Pi04314 and PP1c was obtained via recombinant co-expression in *E. coli* and purification via coupled immobilised metal affinity chromatography (IMAC) and size-exclusion chromatography (SEC). Protein samples were subject to sparse matrix crystallographic screens, and *X*-ray diffraction data were collected from protein crystals at the Diamond Light Source, UK. Structure solution, model refinement, model rebuilding and structure validation were performed as described in the Materials and Methods.

The crystal structure of the Pi04314/PP1c complex reveals Pi04314 contains a single WY domain, which binds to a hydrophobic surface on PP1c (**Figure 1**). From the structure, we identified several hydrophobic residues from the WY domain that are buried at the interface with PP1c, with particular contributions made by Val-100 and Leu-119. These residues make extensive Van der Walls interactions with the backbone and side chains of PP1c residues Ala-196, Gly-215, Met-216, Val-221, Ser-222, and Tyr-223 (**Figure 1 D, E; Figure S1 A**).

**Figure 1.**
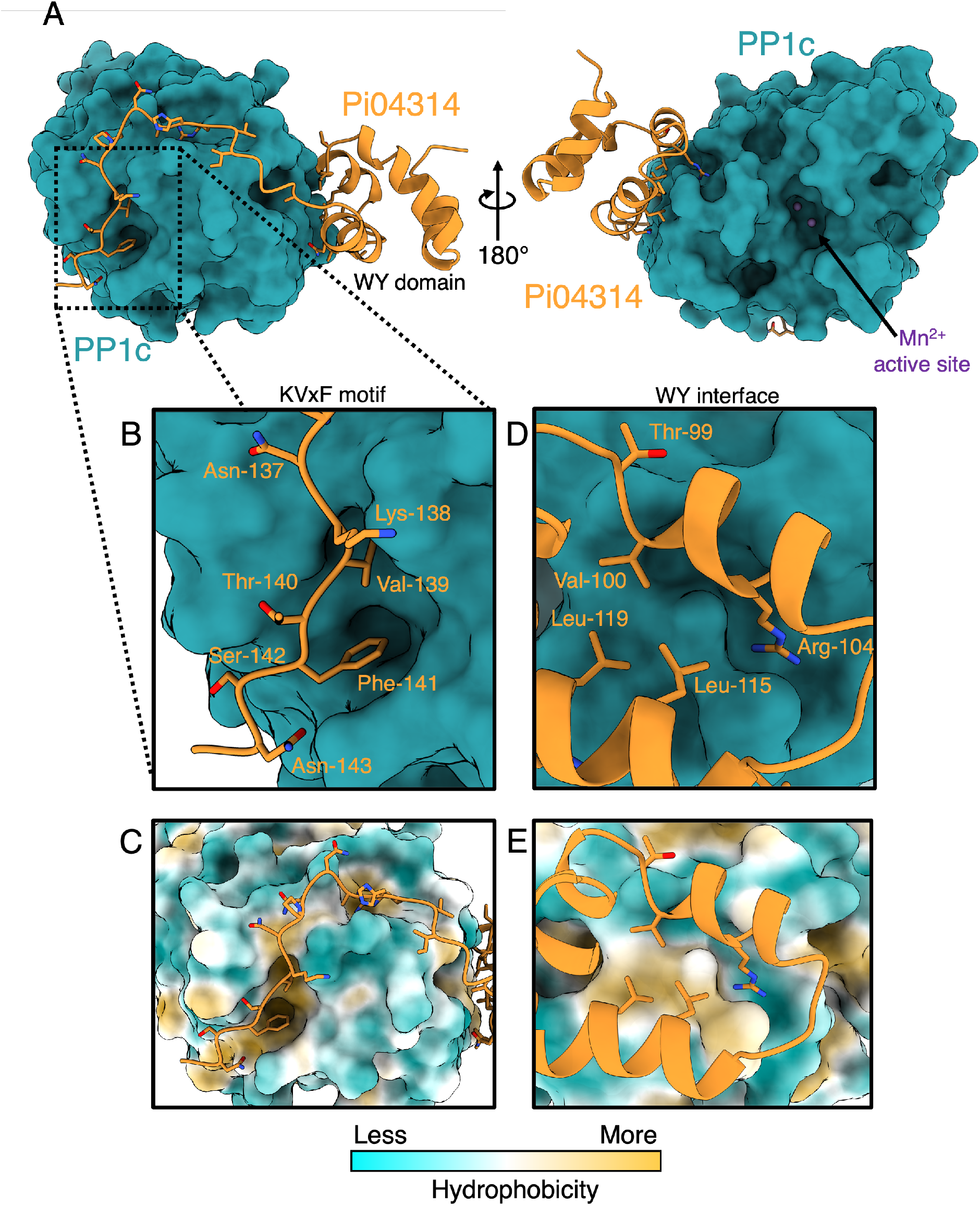
The crystal structure of Pi04314 in complex with PP1c reveals extensive interactions between the WY domain and KVxF motifs and the host target. **A**) The Pi04314 WY domain binds to an anterior surface of PP1c, potentially facilitating the KVxF loop to wrap around the phosphatase surface and position the KVxF motif to bind within the KVxF pocket. **B-C**) The KVxF loop follows a hydrophobic groove across the surface of PP1c and Val-139 and Phe-140 bind to the conserved, hydrophobic KVxF pocket of PP1c. **D-E**) The WY domain of Pi04314 forms an interface via hydrophobic residues (Val-100, Leu-115, Leu-119) flanked by polar/charged residues (Thr-99, Arg-104) to bind an amphipathic surface on PP1c.

The binding of the WY domain likely facilitates the position of the Pi04314 KVxF loop, allowing it to navigate across the surface of PP1c and enable placing of the KVxF motif itself into the canonical phosphatase-binding pocket, as previously established for this motif (Boevink et al. 2016; Terrak et al. 2004). The KVxF loop, to the C-terminus of the WY domain, forms a considerable interface with PP1c comprising amphipathic interactions across nearly every residue of the loop (residues 125-143; **Figure 1 B,C; Figure S1 B**). The conserved Val-139 and Phe-141 residues of the KVxF motif are inserted into the KVxF-binding pocket, formed by PP1c residues Ile-167, Leu-241 Phe-255, Arg-259, Met-288 and Cys-289.

Superimposition of the Pi04314/PP1c complex on the complex of the human PP1B protein and the phosphatase regulator, MYPT1, reveals striking structural similarities (**Figure 2**). Comparison of the two complexes reveals that the Pi04314 WY domain binds to the same surface as the N-terminal arm of MYPT1 (**Figure 2 A, C**), from which a loop region containing the KVxF motif extends. Interestingly, the KVxF loops of MYPT1 and Pi04314 navigate different pathways across the surface of the PP1 proteins, but subsequently converge on the KVxF binding pocket. The conserved valine and phenylalanine residues of the KVxF motifs of both proteins (Val-139 and Phe-141 in Pi04314, and Val-36 and Phe-38 in MYPT1) are bound to the PP1c KVxF pocket in an essentially identical manner (**Figure 2 B)**. The structure of Pi04314 in complex with PP1c determined here demonstrates how pathogen effectors may mimic manipulation of established regulatory surfaces on plant host targets.

**Figure 2.**
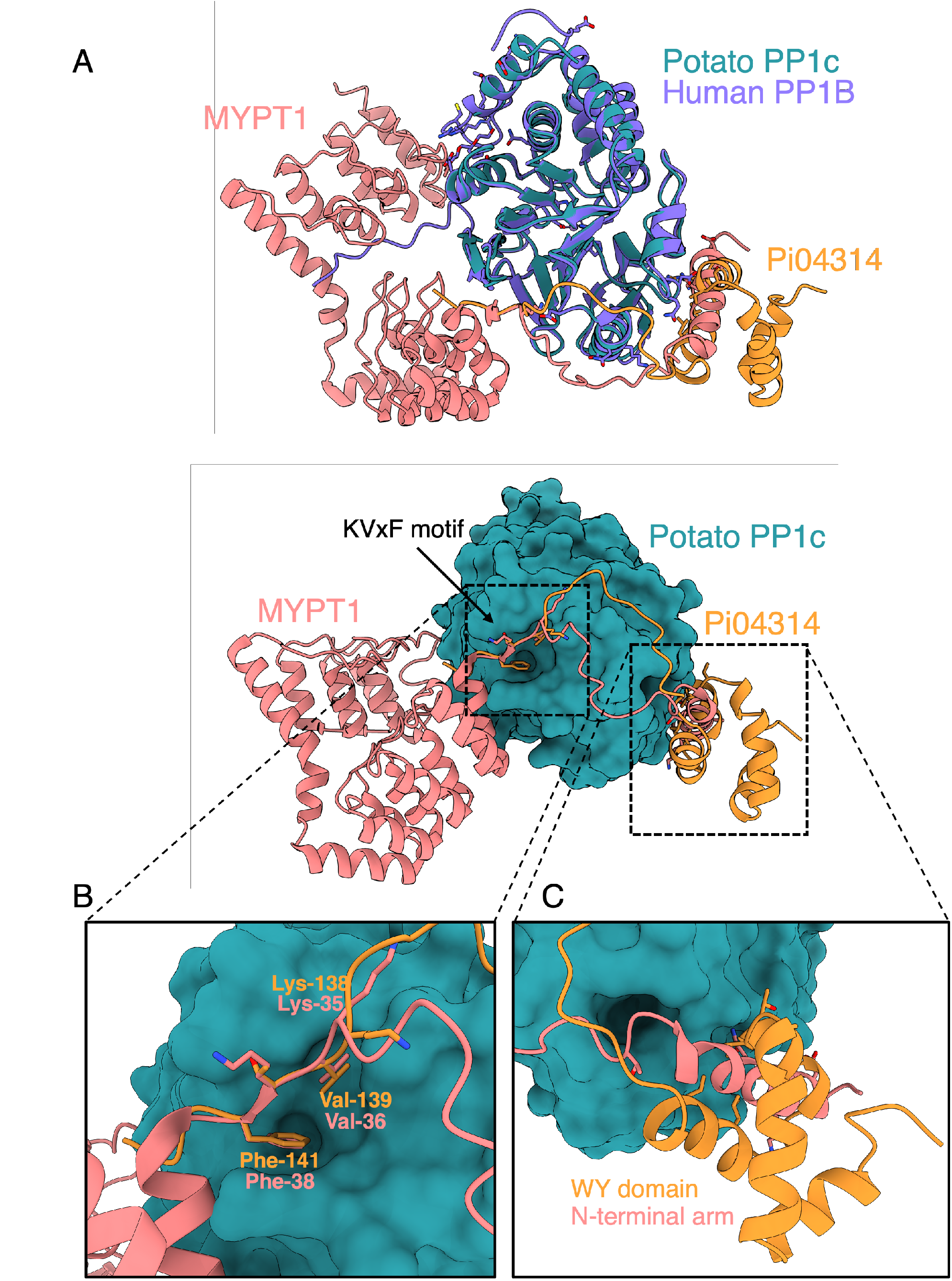
Pi04314 exploits interfaces used by host PP1 regulatory proteins to facilitate interaction. A) Superimposition of the Pi04314/PP1c complex on the structure of the human PP1 regulatory protein MYPT1 bound to PP1B demonstrates Pi04314 occupies two interfaces used by host proteins. B) The KVxF motif Pi04314 binds to PP1c in an essentially identical manner to the KVxF motif of the human PP1 regulator, MYPT1, bound to PP1B. C) The WY domain of Pi04314 interacts the same surface as the N-terminal arm region of the PP1B regulator, MYPT1.

### Biophysical analyses of Pi04314 binding to PP1c suggests contribution from both the WY domain and KVxF motif in complex formation

To assess the importance of the different structural features of Pi04314 on the affinity for PP1c, we performed isothermal titration calorimetry (ITC) assays with recombinantly expressed and purified proteins. For these experiments, we generated structure-guided mutations in Pi04314 to disrupt complex formation with PP1c.

To determine the importance of the WY domain interactions with PP1c, we generated mutants targeting the key residues at the interface as identified in the crystal structure, Val-100 (V100E) and Leu-119 (L119E). We also designed mutants that disrupted KVxF interactions through either mutating the hydrophobic residues to alanine (KATA) or by removing the entire loop, leaving just the WY domain (WY-only). Finally, to help understand the specificity of the Pi04314 WY domain for PP1c, we generated a chimeric effector containing a WY domain from the *P. infestans* effector PexRD54, while maintaining the KVxF loop from Pi04314 (referred to as the WY chimera). This was generated to test potential effects of combining the KVxF loop of Pi04314 with a sequence-divergent but structurally similar WY domain from a different effector.

Isotherms from ITC assays following the injection of WT Pi04314 from the syringe into the cell containing PP1c demonstrated strong binding with dissociation constant (K*_d_*) of 180 nM (**Figure 3**). ITC experiments with Pi04314 mutants each revealed reduction or abolition of binding, with the WY mutants V100E, L199E and the PexRD54 chimera resulting in no detectable interaction with PP1c (**Figure 3**). The KATA and WY-only mutants also significantly reduced the affinity of Pi04314 for PP1c, with K*_d_* values of 1.53 µM and 12.25 µM respectively (**Figure 3**). Our results suggest a significant role for both the WY domain and KVxF motif in PP1c binding in vitro. However, they indicate a substantive role for the WY domain in PP1c interactions, compared to the KVxF motif, as some affinity for PP1c was still observed when the WY domain of Pi04314 remained unchanged, but not when this region was mutated.

**Figure 3.**
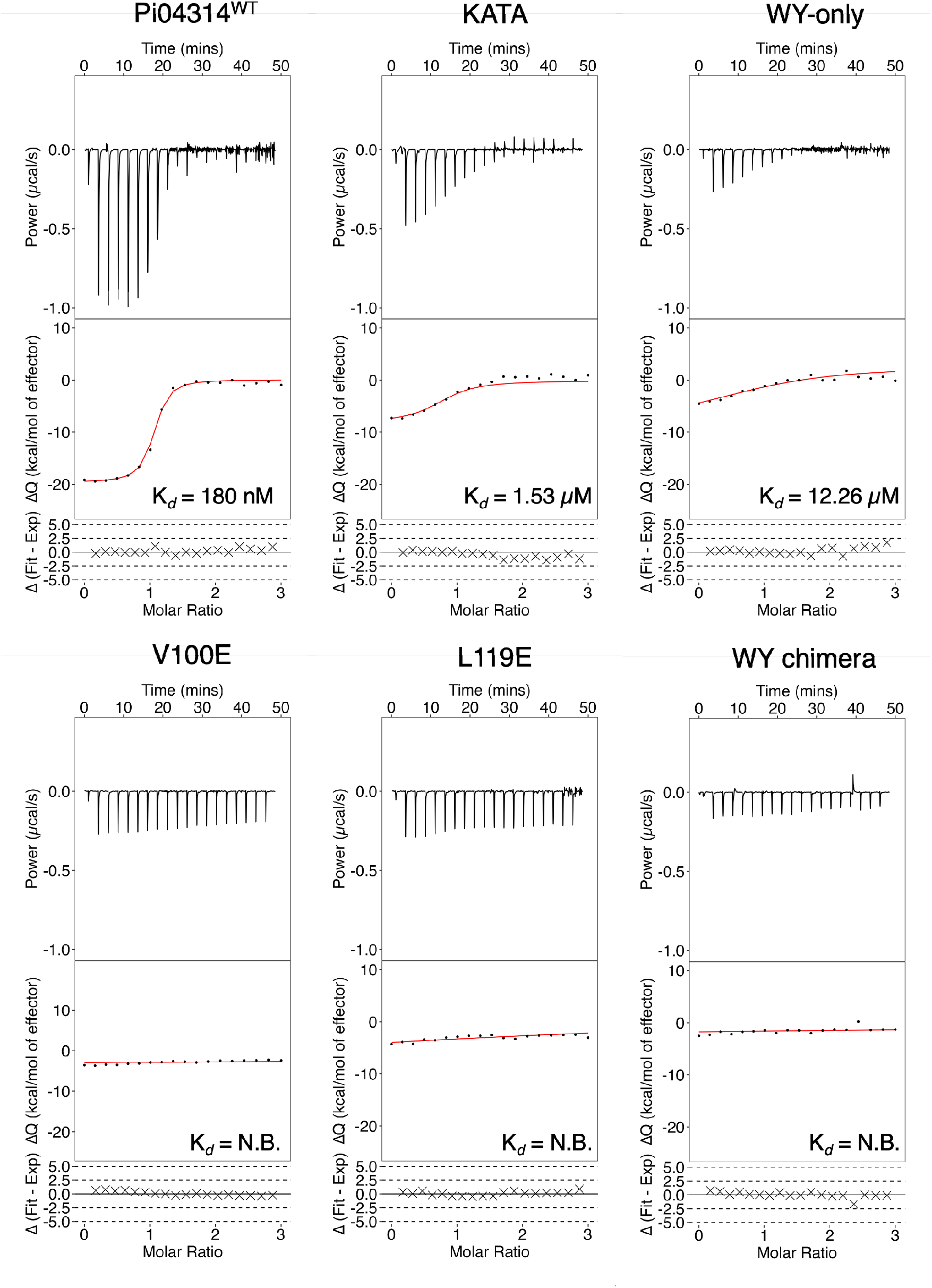
In vitro binding analysis demonstrates mutation of the Pi04314 residues at the WY domain and KVxF interfaces impairs interaction with PP1c in vitro. In vitro binding affinities of Pi04314 and Pi04314 mutants V100E, L199E, KATA, WY-only and WY chimera for PP1c were assessed by isothermal titration calorimetry (ITC). Top panels show heat differences upon injection of the effector into the cell containing PP1c. Middle panel displays integrated heats of injection (•) and the line of best fit (red) to a 1:1 single site model as calculated with the AFFINImeter analysis software. Bottom panel denotes residual differences between experimental and fit data. Wildtype Pi04314 binds PP1c with nanomolar affinity, however mutation to residues of either the the WY domain or the KVxF loop impairs binding, as demonstrated by reduced heat exchange between Pi04314 mutants and PP1c.

### In planta co-immunoprecipitation assays confirm the importance of the WY domain and KVxF motif for association in planta

In addition to the previously established role of KVxF binding to PP1c, our in vitro studies also suggest an important role for the Pi04314 WY domain for interaction with PP1c. To test whether these observations correspond with in planta interaction assays, we performed co-immunoprecipitation (co-IP) using mRFP-tagged WT Pi04314, and the five WY/KVxF mutants, to pulldown GFP tagged PP1c. Constructs were transiently expressed in the heterologous host *Nicotiana benthamiana* via agrobacterium-mediated transformation with tissue harvested 3 days post inoculation (dpi). Immunoblot analysis of samples after incubation with αRFP magnetic beads reveal wildtype Pi04314 was capable of binding PP1c (**Figure 4**). However, none of the Pi04314 mutants V100E, L119, KATA, WY-only or the chimera were able to pulldown PP1c (**Figure 4**). These data correspond with our biophysical analyses in that they demonstrate that mutating either of the WY or KVxF interfaces impacts association of Pi04314 with PP1c. But unlike the in vitro work, mutations in the KVxF motif as well as the WY-region abrogate interaction in planta.

**Figure 4.**
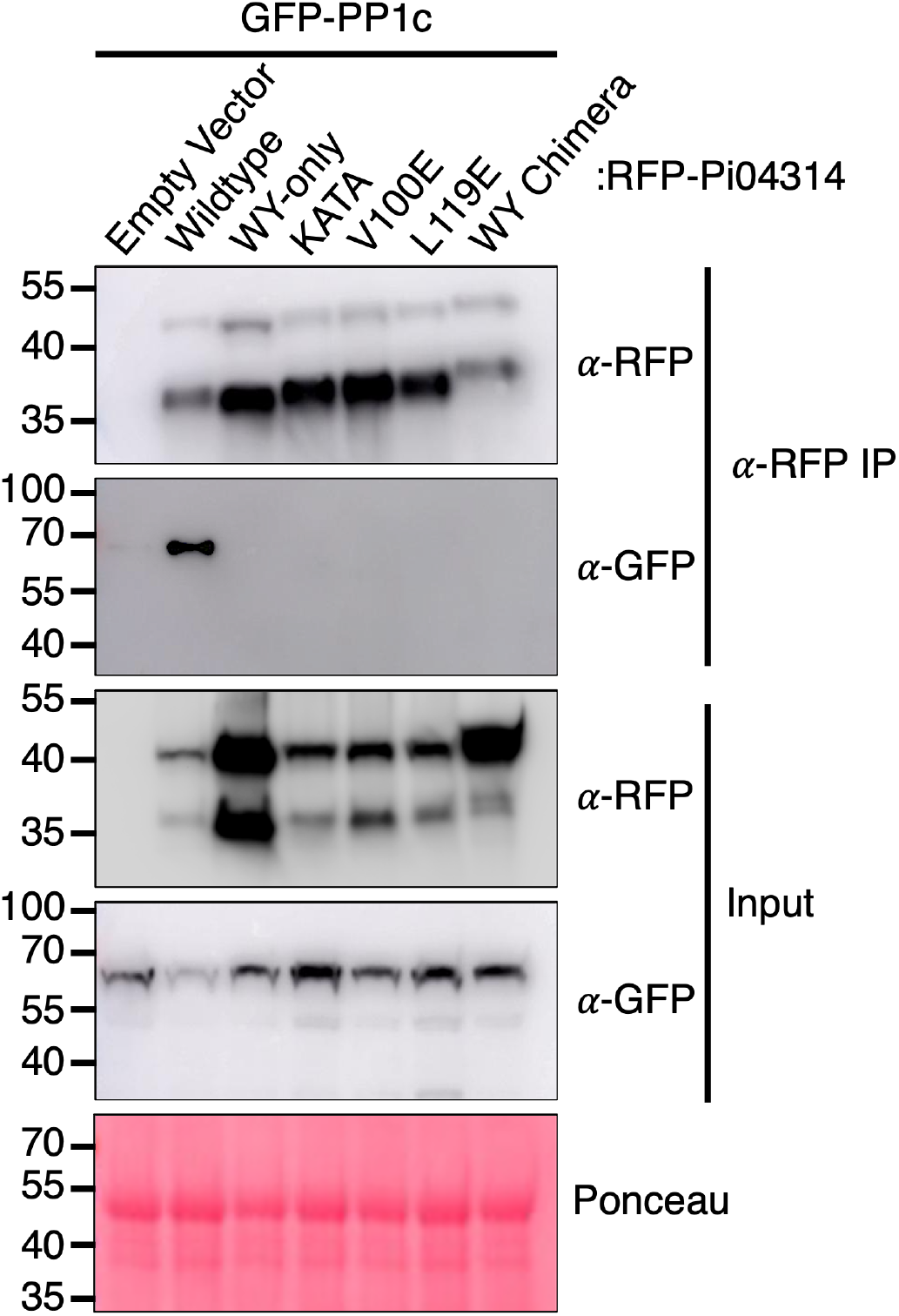
Co-immunoprecipitation experiments reveal Pi04314 interface mutants perturb PP1c binding in planta. Co-immunoprecipitation of GFP-tagged PP1c with Pi04314^WT^ demonstrates a clear association in planta, however interface mutants are unable to pull down PP1c in *N. benthamiana*. Immunoprecipitates obtained with anti-mRFP beads. Immunoblots of IP and input total protein extracts were probed with anti-mRFP and anti-GFP antibodies as appropriate. Total protein loading is shown by Ponceau staining of the input membrane.

### Disruption of Pi04314/PP1c complex formation reduces *P. infestans* virulence in *N. benthamiana* infection assays

Mutation of either the WY domain or the KVxF motif of Pi04314 impacts binding in vitro and in planta. To test whether this disruption of complex formation affects pathogen virulence, we performed pathogenesis assays in *N. benthamiana*.

RFP-tagged wild type (WT) Pi04314 or mutants were transiently expressed on one half of *N. benthamiana* leaves with RFP-EV as a control on the other half. Following *P. infestans* sporangia inoculation, lesions were measured to assess disease development. The half-leaves expressing RFP-tagged WT, WY-only and KATA mutants showed significantly larger lesion sizes (Welch’s two-sample *t* test, n=67, 96 and 86, respectively) compared to the half-leaves expressing free RFP (**Figure 5**). By contrast, no significant differences were observed when comparing V100E, L199E and the WY chimera mutants with RFP-EV control (Welch’s two-sample *t* test, n=83, 68 and 85, respectively, **Figure 5**). These data suggest the integrity of the Pi04314 WY domain is essential for enhancing pathogen virulence in these assays.

**Figure 5.**
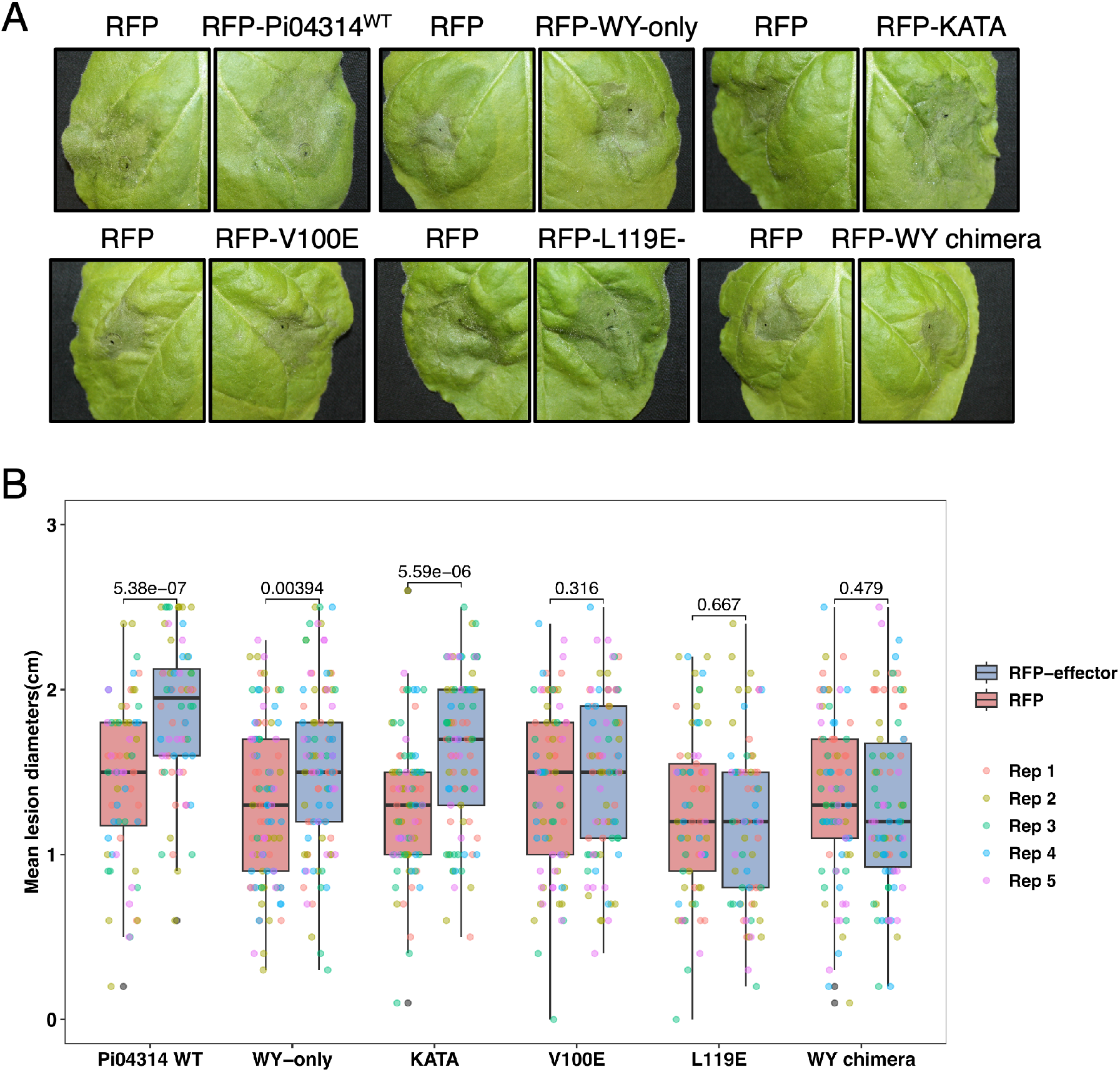
*P. infestans* growth is reduced on *N. benthamiana* leaves expressing the Pi04314 WY domain mutants compared to wild-type or KVxF loop mutants in *N. benthamiana* infection assays. Agrobacterium-mediated transient infection assays with *P. infestans* on *N. benthamiana* tissue transformed with wildtype Pi04314 or mutants. After six days, we observed increased lesion sizes leaves expressing Pi04314^WT^, and the WY-only and KATA mutants, as compared to an mRFP-only negative control. *P. infestans* lesions on leaves expressing Pi04314 WY domain mutants, V100E, L119E and WY chimera, did not show any significant difference in size compared to mRFP control.

### Mutations that disrupt interactions with PP1c reduce the ability of Pi04314 to re-localise the phosphatase from the nucleolus

Previous studies have shown that Pi04314 re-localizes PP1c from the nucleolus to the nucleoplasm (Boevink et al. 2016). Our structural and in planta analyses show that Pi04314 can form a complex with PP1c, suggesting Pi04314 binding to PP1c may function to block entry to or re-localize PP1c from the nucleolus. To test this, we first confirmed the ability of mRFP-Pi04314 to re-localize GFP-tagged PP1c from the nucleolus via confocal microscopy. We observed strong GFP-PP1c signal in the nucleolus in the presence of an mRFP-only control (**Figure 6**). However, co-expression with mRFP-Pi04314 results in reduction of the GFP signal in the nucleolus, suggesting re-localisation of PP1c (**Figure 6**). Next, we tested the ability of Pi04314 mutants with impaired PP1c binding to re-localise PP1c from the nucleolus. All Pi04314 mutants demonstrated a reduced ability to remove PP1c from the nucleolus, with increased GFP intensity in nucleoli when compared to the wildtype Pi04314, showing no significant difference to the mRFP control, with the exception of the WY-only mutant (**Figure 6**). However, the WY-only mutant still demonstrated an impaired ability to re-localise PP1c compared to wildtype (**Figure 6**). This reduced ability of Pi04314 mutants to re-localise PP1c corresponds with reduced binding in our ITC and co-IP assays, indicating the direct interaction between Pi04314 with PP1c is important for this effector function.

**Figure 6.**
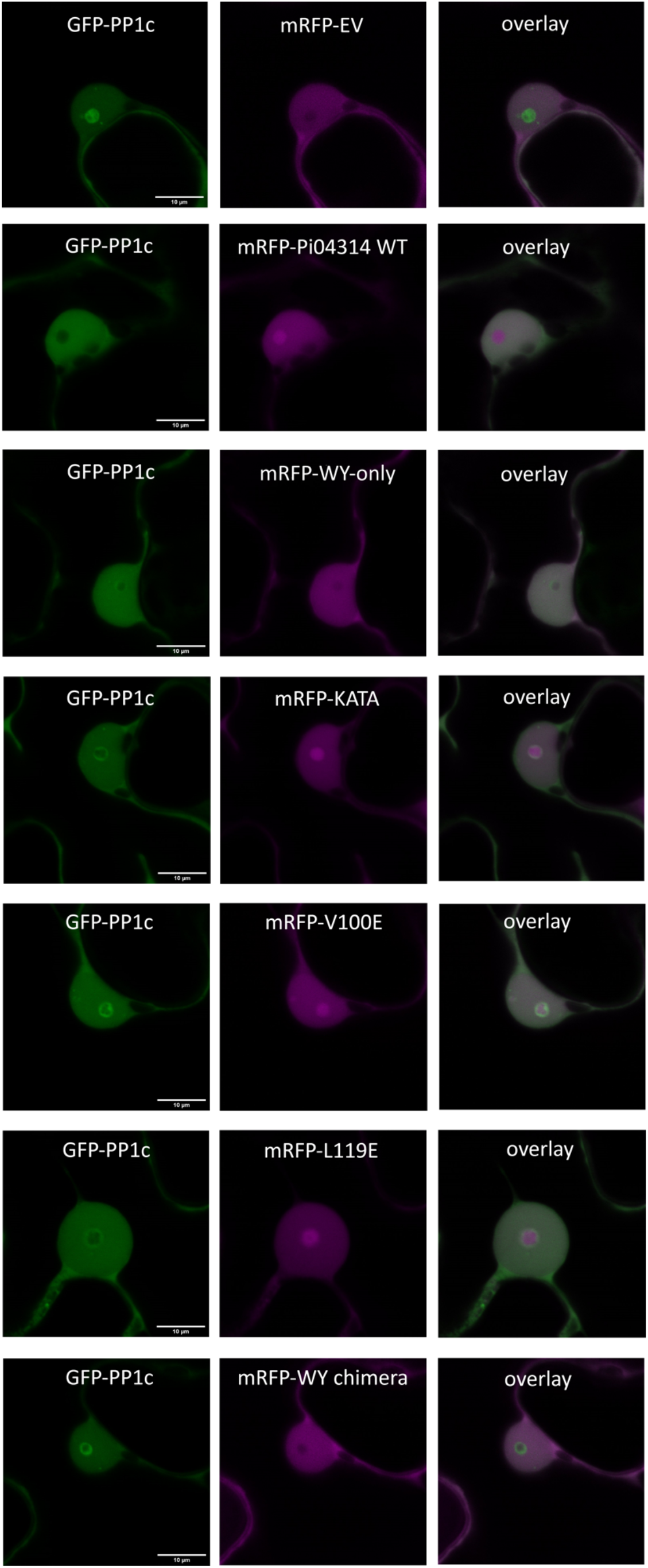
Pi04314 mutants show reduced ability to re-localise PP1c from the nucleolus. Single optical sections through *N. benthamiana* nuclei demonstrate Pi04314 mutants with impaired PP1c binding have reduced ability to re-localise PP1c from the nucleolus compared to wildtype Pi04314. All mutants demonstrated no significant difference in nucleolar GFP intensity compared to an mRFP empty vector control, with the exception of the WY-only mutant, which could re-localise some PP1c, but not to the same levels as Pi04314^WT^. Scale bar represents 10 µM. Images sets are representative of 90 nuclei for each protein.

## Discussion

Effectors play crucial roles in the successful colonisation of plants by plant pathogens and demonstrate a wide range of protein folds and functions. In this study, we investigated the function of the Pi04314 effector from *P. infestans*, the causal agent of potato blight, and its role in promoting pathogen infection through direct interaction with a host target phosphatase, PP1c. Structural analysis of the Pi04314/PP1c complex provides valuable insights into the binding mechanism between pathogen and host proteins, highlighting the significance of the WY domain in facilitating target binding. In addition, the C-terminal KVxF motif of Pi04314 engages with the canonical phosphatase-binding site on the surface of PP1c.

The crystal structure of the Pi04314/PP1c complex shows that Pi04314 comprises a single WY domain, decorated with a C-terminal loop containing a KVxF motif. As a well-established motif used for regulation and interaction with PP1 proteins (Terrak et al. 2004), we expected the KVxF motif to have a role in PP1c binding through similar mechanisms to host regulatory proteins, as demonstrated previously (Boevink et al. 2016). However, the crystal structure also revealed the WY domain to bind to a conserved surface on the PP1c protein, and through structural comparison with the human MYPT1 protein bound to PP1B, we could visualise how both the Pi04314 WY domain and KVxF loop could facilitate the binding to PP1c regulatory surfaces. MYPT1 serves as a crucial regulatory subunit of PP1 proteins in mammalian systems, playing a role in guiding PP1 proteins towards specific targets for dephosphorylation (Terrak et al., 2004a; Scotto-Lavino et al., 2010).

While it remains unknown whether Pi04314 functions similarly to MYPT1 in directing PP1c to specific dephosphorylation targets, it is evident that this effector utilizes overlapping regulatory interfaces employed by signalling partners of PP1c to direct PP1c towards compartments in healthy systems. Redirection of PP1c away from the nucleolus by Pi04314 contributes to successful *P. infestans* infection. Intriguingly, there is no structural evidence to suggest Pi04314 binding to PP1c effects PP1c enzymatic function. Pi04314 binding occurs away from the Mn^2+^ active site of the PP1c (**Figure 1 A**), with no apparent allosteric disruption of Mn^2+^ ion coordination or significant structural changes to the local area around the catalytic residues, as compared to the active human PP1B in complex with regulator MYPT1. This suggests PP1c likely remains an active phosphatase when bound by Pi04314, as suggested by previous experiments (Boevink et al. 2016). It is possible Pi04314 may manipulate PP1c to dephosphorylate specific host targets, favouring *P. infestans* infection.

Our biophysical and in planta analyses support the hypothesis that re-localization of PP1c by Pi04314 occurs through direct interaction mediated by the WY domain and the KVxF loop. Mutations introduced to Pi04314, which hinder its binding with PP1c, also demonstrate a diminished capacity to re-localize PP1c. This diminished re-localisation is consistently associated with reduced pathogen success in our infection assays.

Interestingly, our findings revealed that mutations affecting the binding of the WY domain to PP1c had a more pronounced impact on the function of the effector compared to mutations targeting the KVxF motif. In isothermal titration calorimetry (ITC) assays, Pi04314 mutants with a wild-type WY domain still exhibited a reduced but detectable binding to PP1c, unlike the mutants targeting residues of the WY domain. In particular, the WY-only mutant, lacking the entire KVxF motif loop, still supported promotion of pathogen growth in infection assays compared to a negative control, along with a partial capability to re-localize PP1c away from the nucleolus. These findings suggest that the WY domain of Pi04314 has an important role in facilitating the interaction with PP1c, while the KVxF motif may act to strengthen the interaction between the host target and the effector. Previously, a quadruple alanine mutant, replacing the KVxF motif entirely, did not re-localise PP1c or support enhanced pathogen growth (Boevink et al. 2016). It is possible that these different mutants support subtly different phenotypes in these assays.

Previous studies of another *P. infestans* WY/LWY effector, PSR2, demonstrated the importance of WY domains in directly binding to its host target Protein Phosphatase 2A (PP2A) (He et al. 2019; Li et al. 2023). Studies of mammalian PP2A and PP1c proteins have shown they share several targets in common, suggesting functional redundancy (Wlodarchak and Xing 2016; Gil and Vagnarelli 2019; Sontag et al. 2008). However, less is known about the overlap between PP1c and PP2A functions in the context of plant signalling pathways. The work presented here, in combination with previous studies of PSR2, implies convergence of these two WY effectors on similar signalling pathways, with both functioning to manipulate host phosphatases. However, these shared functions are mediated through distinct mechanisms. While Pi04314 utilises a single WY domain with an attached KVxF motif, the PSR2 effector is a large multi-domain effector consisting of a WY domain followed by several tandem LWY domains which function to mimic subunits of the PP2A holoenzyme (Li et al. 2023). Comparison of these two effectors suggests convergent evolution of WY domain effectors to target similar pathways, but in mechanistically distinct manners.

Our study highlights the capacity of WY effectors to bind and manipulate host targets by using conserved regulatory surfaces. The WY domain and KVxF motif of Pi04314 play vital roles in effector function by facilitating interactions with PP1c, enabling the re-localization of the host target, and enhancing pathogen virulence in plant assays. Further investigations should aim to elucidate whether Pi04314 re-localizes PP1c to specific host proteins to mediate their dephosphorylation, therefore modulating signalling pathways in a targeted manner. Understanding these mechanisms would be a valuable direction for future research in detailing the functions of effectors in plant-pathogen interactions.

## Materials and Methods

### Gene cloning – recombinant expression in *Escherichia coli*

The gene encoding the PP1c^1-293^ protein (herein simply referred to as PP1c) was cloned into the pOPIN-SC3 vector (OPPF) via In-Fusion Cloning (Takara Bio), which provides a cleavable N-terminal 6xHIS-SUMO tag for purification and solubility. For Pi04314, the N-terminal before the RxLR motif was removed, leaving the WY and C-terminal KVxF loop for expression (residues 68 – 254; herein referred to as Pi04314). Pi04314 was also cloned into pOPIN-S3C via In-Fusion Cloning. For co-expression and crystallisation, an untagged version of PP1c was required. To create this construct, PP1c^1-293^ was cloned into pOPIN-A via In-Fusion Cloning.

The Pi04314 mutants V100E, L119E and KATA were generated via site-directed mutagenesis of the Pi04314 in the pOPIN-S3C vector. For the WY-only, primers were designed to truncate the C-terminal region at residue Arg-131. To generate the WY chimera, a gblock containing the WY domain of PexRd54 of corresponding length to the Pi04314 WY domain (residues 68 – 131) and the C-terminal KVxF loop of Pi04314 (residues 132 – 154) was synthesized (IDT DNA) with In-Fusion Cloning sites to allow cloning into pOPIN-S3C.

### Protein expression and purification from *E. coli*

The pOPIN-S3C expression vector containing PP1c was transformed into *E. coli* BL21 (DE3) cells. Using an overnight culture for inoculum, 8 L of BL21 DE3 cells were grown in LB media at 30°C to an OD_600_ of 0.6 – 0.8 before the temperature was reduced to 18°C for expression via the addition of 1 mM IPTG. For Pi04314, expression vectors were transformed into BL21 arabinose inducible (AI) cells. Similar to PP1c, 8 L of BL21 AI cells were grown in LB media at 30°C to an OD_600_ of 0.6 – 0.8 before the temperature was reduced to 18°C for expression via the addition of 0.2 % v/v l-arabinose.

After growth, cells were harvested via centrifugation at 5000 *g* for 10 mins and resuspended in lysis buffer (50 mM HEPS pH 8.0, 500 NaCl, 30 mM imidazole. 50 mM glycine and 5 % v/v glycerol. Cells were lysed via sonication and then subjugated to centrifugation at 45,000 *g* for 20 mins to clarify the lysate. Proteins were purified from the cell lysate via Ni^2+^ immobilised metal affinity chromatography (IMAC) couple with size exclusion chromatography (SEC). The 6xHIS-SUMO tags were removed from the PP1c and Pi04314 proteins via overnight proteolytic cleavage with 3C protease at 4C, followed by a reverse IMAC step before a final round of SEC using a buffer of 10 mM HEPES pH 8.0, 150 mM NaCl and 1 mM TCEP. Proteins were vitrified in liquid nitrogen before storage at −80°C.

For production of the Pi04314 / PP1c complex, the pOPIN-S3C-Pi04314 and pOPIN-A-PP1c (untagged) vectors were co-expressed in *E. coli* BL21 AI. 8 L of BL21 AI cells were grown in LB media at 30°C to an OD_600_ of 0.6 – 0.8 before the temperature was reduced to 18°C for expression via the addition of 0.2 % v/v l-arabinose. The complex was purified under the same conditions as the individual proteins.

### Crystallisation, x-ray data collection, structure solution and refinement

The Pi04314 / PP1c complex was concentrated to 10 mg/mL in SEC buffer (10 mM HEPES pH 8.0, 150 mM NaCl and 1 mM TCEP) for crystallisation. Sitting drop vapour diffusion crystallisation trials were performed in 96-well plates using an Oryx Nano robot (Douglas Instruments). Crystallisation plates were incubated at 20°C. Crystals of the Pi04314 / PP1c appeared in a condition consisting of 0.2 M sodium fluoride, 0.1 M BIS-TRIS propose pH 8.5 and 20 % PEG3350. Crystals were harvested and vitrified in liquid nitrogen prior to shipping to the Diamond Light Source, Oxford.

Crystals of the Pi04314 / PP1c complex diffracted to 2.1 Å and *X*-ray datasets were collected at the i04 beamline, Diamond Light Source under proposal mx18565. Data were processed using the xia2 pipeline and AIMLESS as implemented in CCP4i2 (Winn et al. 2011). The structure of the RepoMan-PP1g proteins (PDB ID: 5inb) was used as a search model for PP1c, to solve the structure via molecular replacement with PHASER (McCoy et al. 2007), which placed two copies of the phosphatase in the asymmetric unit. Then, automated model building with BUCCANEER was used to de novo build to molecules of the Pi04314 before refinement through iterative cycles of model building in COOT, geometry correction in ISOLDE and refinement in REFMAC (Emsley and Cowtan 2004; Croll 2018; Murshudov et al. 1997; Cowtan 2006). Structure geometry was validated using MOLPROBITY (Chen et al. 2010). Analyses of the protein interfaces were performed with ChimeraX (Pettersen et al. 2021). The final structure and the *X*-ray diffraction data used to derive this have been deposited at the Protein Data Bank with the accession number 8PQ7.

### Isothermal titration calorimetry in vitro binding analysis

All calorimetry experiments were recorded using a MicroCal PEAQ-ITC (Malvern, UK). Binding assays were performed in triplicate at 25°C in SEC buffer (10 mM HEPES pH 8.0, 150 mM NaCl and 1 mM TCEP). The calorimetric cell was filled with 20 µM PP1c and titrated with 300 µM Pi04314. For each ITC run, a single 0.5 µL injection of Pi04314 was followed by 19 subsequent 2 µL injections. Injections were made at 120 second intervals, with a stirring speed of 750 rpm. This was repeated for each of the Pi04314 mutants: V100E, L199E, WY-only, KATA and WY chimera. As informed by the crystal structure, data were fit to a 1:1 binding model and processed with the AFFINImeter ITC analysis software (Piñeiro et al. 2019). Processed data were visualised using the ggplot2 R library in R 4.2.2 (Wickham 2016).

### In planta co-immunoprecipitation

For in planta co-immunoprecipitation (co-IP), PP1c and Pi04314 were cloned into the pICH47742 plant binary expression vector via GoldenGate cloning with *Bsa1,* GFP and mRFP tags, respectively. PP1c and Pi04314 were transiently expressed in planta via agrobacterium-mediated transformation of four week old *N. benthamiana* plants with *A. tumefaciens* strain GV3101 (C58 (Rif*^R^*) Ti pMP90 (pTiC58DT-DNA) (Gent*^R^*) Nopaline (pSoup-Tet*^R^*). *A. tumefacians* carrying PP1c and Pi04314 were infiltrated at an OD_600_ of 0.5 in agrobacterium infiltration medium (10 mM Mg Cl_2_, 10 mM 2-(N-morpholine)-ethanesulfonic acid (MES), pH 5.6) with the addition of 150 µM acetosyringone. Tissue from several different infiltrations were used for different replicates.

Leaf tissue was collected three days post infiltration (dpi) and frozen in liquid nitrogen before processing. Leaf tissue was collected 3 days post infiltrations (dpi) and frozen in liquid nitrogen before processing. Samples were ground to a fine powder in liquid nitrogen using a mortar and pestle before being mixed with two times weight/volume ice-cold Co-IP extraction buffer (50 mM HEPES pH 8.0, 150 mM NaCl, 1 mM EDTA, 20 % v/v glycerol, 2 % w/v PVPP, 10 mM DTT, 1 x cOmplete protease inhibitor tablet per 50 mL (Roche), 0.1 % Tween20). Sample lysates were centrifuged at 15,000 *g* at 4°C for 20 min to clarify. SDS-PAGE/immunoblot analysis was used to identify proteins in the sample with use of anti-GFP antibody (Cohesion Biosciences) and anti-mRFP antibody (Abcam) for PP1c and Pi04314 proteins, respectively.

For immunoprecipitation, 600 µL of plant lysate was incubated with 30 µL of anti-mRFP magnetic beads (Chromotek) in a rotary mixer for 1 hrs at 4°C. The mRFP beads were separated from the supernatant with use of a magnetic rack to allow for the removal of the supernatant. The beads were then washed three times with 1 mL of IP buffer (50 mM HEPES pH 8.0, 150 mM NaCl, 1 mM EDTA, 20 % v/v glycerol, Tween20). After washing, 30 µL of LDS Runblue sample buffer was added to the mRFP beads and incubated for 5 min at 95°C. The beads were then applied to a magnetic rack and the supernatant was loaded to SDS-PAGE gels and subsequently used for immunoblot blot analysis. PVDF membranes were probed with anti-mRFP and anti-GFP antibodies to detect Pi04314 proteins and PP1c, respectively. mRFP immunoprecipitation and subsequent immunoblot analyses were performed in duplicate using tissue from different agrobacterium infiltrations.

### Agrobacterium-Mediated Transient Infection Assays

Agrobacterium expressing different RFP-tagged Pi04314 constructs and RFP-EV control were grown in yeast extract and beef media (YEB) with antibiotics and shaking at 28 °C overnight. Cultures were centrifuged at 3900 rpm to obtain the bacteria pellet and resuspended in 1×MES buffer (10 mM MES, 10 mM MgCl_2_ and 200 mM acetosyringone). Optical densities (ODs) were adjusted by MES buffer to 0.1 at 600 nm and left in dark for at least 1 hr ahead of infiltration. Agrobacterium suspensions containing RFP-effector or RFP-EV were then infiltrated into either half of *N. benthamiana* leaves. Both infiltrated leaf halves were drop inoculated with 10 μl *P. infestans* sporangia (50,000 sporangia/ml) one day after infiltration. Lesion sizes were measured at 6 days post-inoculation (dpi). Fifteen to eighteen detached leaves from six individual plants were used and 5 independent replicates of this were performed.

### Confocal microscopy and protein localisation assays

Agrobacterium expressing GFP-PP1c was co-infiltrated with different mRFP tagged effectors or mRFP empty vector at a low OD_600_(0.01-0.05). *N. benthamiana* leaf cells were imaged no later than 2 days after agroinfiltration using Leica TCS SP2 AOBS, Ziess 710. GFP was excited by the 488-nm line of an argon laser, and emissions were detected between 500 and 530 nm. mRFP was excited with a 561-nm line from a diode laser, and their emissions were collected between 580 and 610 nm or 600 and 630 nm, respectively. The pinhole was set at 1 airy unit for the longest wavelength of light being used. To quantify the relative GFP intensity in the nucleolus, the nucleolar to nucleoplasmic ratio of fluorescence intensity was chosen as a measure of re-localisation to account for variation of protein expression levels from cell to cell, 30 images of nuclei were taken for each of 3 replicates. Images were processed and quantified by ImageJ.

## Acknowledgements

We thank the Diamond Light Source, UK (beamline i04 under proposal mx18565) for access to *X*-ray data collection facilities and Dave Lawson from the JIC Protein Crystallography Platform for expert technical assistance during data collection. We also thank Clare Stevenson and Julia Mundy from the JIC Biophysical Analysis platform for their help with ITC.

---

**Supplemental Figure S1.**
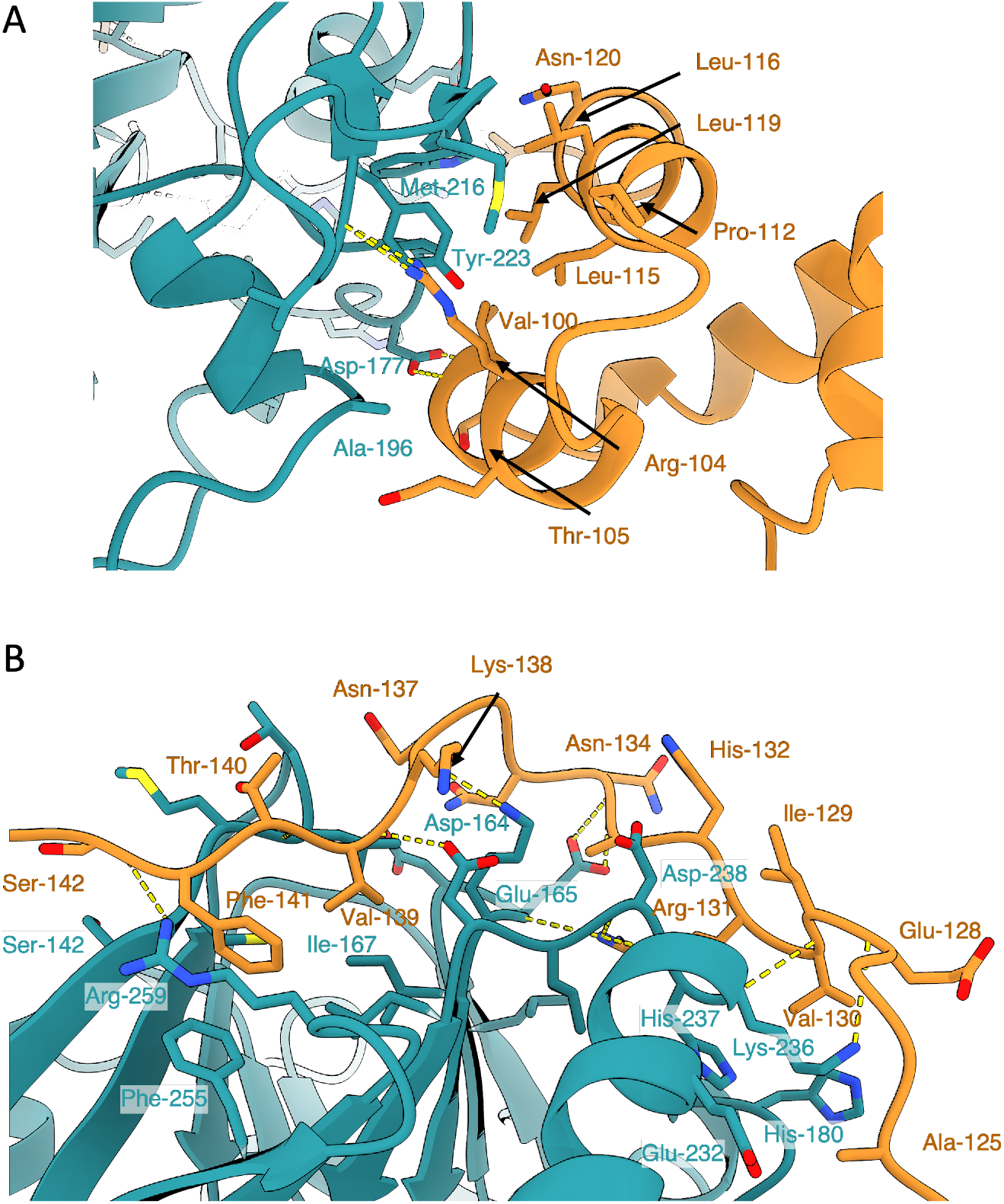
Atomic contacts and hydrogen bonding at the WY and KVxF interfaces. A) Interactions between the residues of the Pi04314 WY domain and PP1c. B) Interactions between the residues of the KVxF loop and PP1c. Yellow dashed lies denote hydrogen bonds.

**Supplemental Figure S2.**
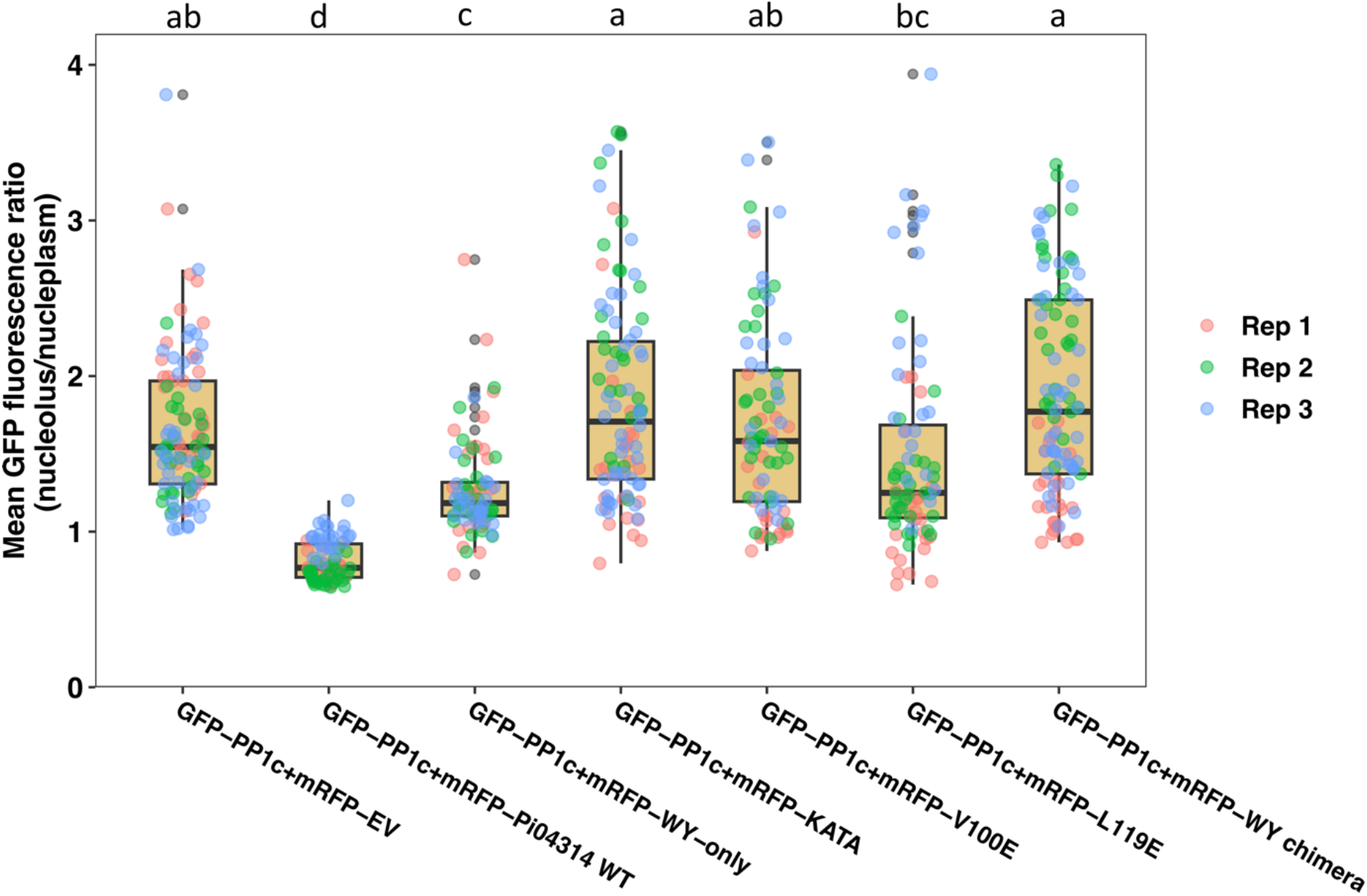
Quantification of GFP intensity in the nucleolus in the presence of wildtype Pi04314 and mutants. Boxplots of average ratios mean ratios of nucleolar to nucleoplasmic GFP fluorescence from the PP1c expressed alone or with Pi04314^WT^ and mutants, demonstrating significant difference in the ability of Pi04314 mutants to re-localise PP1c. Averages were obtained from images of X nuclei for each sample. Error bar represent standard error and the graph represents the combined data from X biological replicates. Letters on the graphs denote statistically significant differences (one-way ANOVA with Tukey’s HSD post-hoc test)

**Supplemental Table S1.** X-ray data collection and refinement statistic for the crystal structure of PP1c / Pi04314.

|  | Pi04314/PP1c |
| --- | --- |
| **Data collection statistics** |  |
| Wavelength (Å) | 0.91188 |
| Space group | P1 |
| Cell dimensions (*a*, *b*, *c*, $\alpha$, $\beta$, $\gamma$) | 55.49 Å, 57.04 Å, 59.02 Å, 84.35°, 86.49°, 89.85° |
| Resolution (Å)* | 58.62 - 2.10 (2.16 - 2.10) |
| $R_{\text{meas}}$ (%) | 9.7 (52.7) |
| $I/\sigma I$ | 9.2 (1.8) |
| Completeness (%) |  |
| Overall | 97.6 (96.6) |
| Anomalous | 93.2 (92.2) |
| Unique reflections | 40927 (3300) |
| Redundancy |  |
| Overall | 3.4 (3.2) |
| Anomalous | 1.7 (1.6) |
| $CC^{(1/2)}$ | 0.997 (0.840) |
| **Refinement and model statistics** |  |
| Resolution (Å) | 58.62 - 2.10 (2.16 - 2.10) |
| $R_{\text{work}}/R_{\text{free}}$ (%) | 20.2 / 25.1 |
| No. atoms |  |
| Protein | 5889 |
| Ligand | 12 |
| B-factors |  |
| Protein | 24.32 |
| Ligand | 45.16 |
| R.m.s deviations |  |
| Bond lengths (Å) | 0.0086 |
| Bond angles (°) | 1.661 |
| Ramachandran plot (%)** |  |
| Favoured | 93.49 |
| Allowed | 5.96 |
| Outliers | 0.55 |
| MolProbity Score | 2.38 |

\*The highest resolution shell is shown in parenthesis.
\*\*As calculated by MolProbity

